# The hyper-attenuated RVFV 40Fp8 strain can be safely administered to pregnant ewes to protect them from a virulent challenge

**DOI:** 10.1101/2025.01.22.634310

**Authors:** Belen Borrego, Celia Alonso, Sandra Moreno, Eva Calvo-Pinilla, Gema Lorenzo, Pedro J. Sánchez-Cordón, Alejandro Brun

## Abstract

In the present study, we evaluated the immunogenicity, safety and protective efficacy of the attenuated RVFV-40Fp8 strain in natural hosts (non-pregnant ewes) and in a highly susceptible host infection model such as pregnant ewes in the first third of pregnancy. Our results confirm the immunogenicity of 40Fp8 administration in non-pregnant and pregnant ewes, as well as the absence of foetal damage even after a high-dose vaccination regime in pregnant ewes. In addition, the ewes and their foetuses were protected against a virulent RVFV-56/74 strain challenge, as shown by comparative histopathological evaluation of tissue samples from vaccinated and non-vaccinated pregnant ewes. These results confirm the potential use of 40Fp8 as a RVF live-attenuated vaccine candidate and pave the way for further clinical developments.

## Introduction

Rift Valley fever (RVF) is a zoonotic arboviral disease that causes abortions and deaths in livestock (cattle, sheep, goats and camels), and in wild ruminants [1]. The disease is caused by the Rift Valley fever virus (RVFV), a well-known phlebovirus (*class Bunyaviricetes, Order Hareavirales, family Phenuiviridae*), that is transmitted and perpetuated in nature by competent mosquito vectors of the *Aedes* and *Culex* genera [2–5]. RVFV virions display an icosahedral symmetry [6, 7] and contain at least three ssRNA genome segments of different size and negative (large and medium segments) or ambisense (small segment) polarities. The large (L) segment encodes for the virion associated RNA polymerase (RdRp). The medium (M) segment contains five in-frame AUG codons that are alternatively used for the synthesis of two major structural glycoproteins, Gn and Gc, and at least two accessory proteins, NSm, a 13-14-kDa cytosolic protein, and a 78-kDa glycoprotein equivalent to a fused NSm-Gn ORF [8]. The small (S) segment encodes the nucleocapsid (N) protein and the accessory protein, NSs in opposite orientation separated by an intergenic region (IGR). The disease is endemic in Africa and southern regions of the Arabian Peninsula. To ameliorate the impact of the disease in rural and peri urban communities, vaccination of susceptible livestock has been proposed as a “One-Health” approach to prevent disease outbreaks [9–11].

Live attenuated vaccines (LAVs) are excellent inducers of protective and memory immune responses. Their success is largely based on limited replication in the infected host and the induction of a functional immune response. In some cases, the use of LAVs may not be advisable due to potential adverse effects, particularly in immunosuppressed individuals or during pregnancy, either because of impaired immune responses that may not control virus replication, or because of the greater susceptibility of developing foetal tissues to viral invasion. Several live attenuated vaccine candidates have been reported in the past for RVFV, some of which are commonly used in African countries as a means of disease control or are in early clinical development [12, 13]. The guidelines for a LAV development for RVF require, among other several considerations, to ensure safety (see (https://www.who.int/publications/m/item/who-target-product-profiles-for-rift-valley-fever-virus-vaccines), therefore a careful determination of residual virulence should be carried out first.

In a previous study, we obtained and characterized a novel mutagenized RVFV variant strain (40Fp8) that showed a hyper-attenuated phenotype “in vivo” [12]. In murine infection models, the inoculation of the attenuated 40Fp8 strain induced strong protective immunity, thereby making this virus an attractive candidate for vaccination. It was noteworthy that the inoculation of immunodeficient A129 IFNAR^-/-^ transgenic mice, considered as a highly susceptible infection model, with 40Fp8 was not pathogenic, even after intranasal inoculation, while other attenuated vaccine candidates such as rMP12 or NSs-deleted virus induced clear pathological damage [13]. This higher level of attenuation displayed by 40Fp8 in the immunodeficient murine model was therefore an important asset to exploit its suitability as a safer LAV.

In this study, we aimed to test the immunogenicity, safety and the protective efficacy of the RVFV-40Fp8 strain against a virulent challenge, using non-pregnant ewes together with pregnant ewes in the first third of gestation, the latter considered as an animal model extremely susceptible to RVFV infection, by monitoring infectivity in ewes, reproductive tissues and foetuses. The results shown here confirm the immunogenicity of the vaccine candidate in ewes as well as its safety and protective capability in this target species.

## Methodology

### Ethics statement

Procedures involving the use of animals were conducted in accordance with Spanish regulations on animal experimentation (RD 53/2013) and European regulations (EU directive 2010/63/EU) on the protection of animals used for scientific purposes. The animal research ethics committee (Consejo Superior de Investigaciones Científicas-CSIC) and the regional authorities of the Community of Madrid (permit number PROEX/22/06.79) approved the experimental procedures.

### Cells and viruses

Vero and/or Vero E6 cell lines (ATCC #CCL-81 and #CRL-1586) were grown and subcultured in DMEM supplemented with 2-5% foetal bovine serum under conventional conditions of temperature (37°C) and humidity (95%) with a 5% CO2 atmosphere. The RVFV-40Fp8 virus was previously described [12, 13]. The original isolate was propagated upon low moi (0.02) Vero cell inoculation [13]. The supernatant was clarified, titrated, and used as inoculum for experiments 1 and 2 (see below). For pregnant sheep inoculation (experiments 3 and 4), a larger stock of 40Fp8 was produced by amplification of a plaque-purified clone from the original isolate obtained after 8 passages in Vero cells in the presence of 40 µM of favipiravir [12, 14]. Briefly, several biological clones picked up from infected Vero cells were first grown in the presence of the drug, supernatants titrated and further amplified at a moi of 0.1 in the absence of the drug. 72 hpi supernatants obtained were collected, clarified, titrated, and conserved in aliquots at-80°C. A stock designated as c1/2022 was finally selected as inoculum for the animal experiments. The genetic stability of c1/2022 was confirmed by Sanger sequencing as described [15]. Challenge virus was isolated from an experimentally infected sheep that succumbed to infection with the virulent RVFV 56/74 South African origin isolate [16, 17]. This virus was further grown and propagated in C6/36 mosquito cells (CISA repository) as previously described [14, 18].

### Animal study design

Four experiments were conducted to confirm the immunogenicity, safety and protective efficacy of the live attenuated 40Fp8 prototype vaccine against RVFV in non-pregnant ewes and pregnant ewes of different Spanish breeds, as the ovine species is a natural host of the virus and highly susceptible to the disease (**figure 1**). To assess immunogenicity after 40Fp8 inoculation, two pilot experiments were performed using *Churra* ewes (experiments 1 & 2). Safety was further tested in pregnant *Manchegan* ewes (experiment 3) while, to test protective efficacy upon virus exposure, *Castilian* pregnant ewes vaccinated with 40Fp8 were challenged with the virulent 56/74 isolate (experiment 4). Animals were supplied by commercial farms with high sanitary standards. All animals underwent a 7-day acclimatization period in the BSL-3 animal facility of the Animal Health Research Center (CISA-INIA/CSIC) before starting the experiments.

**Fig. 1.**
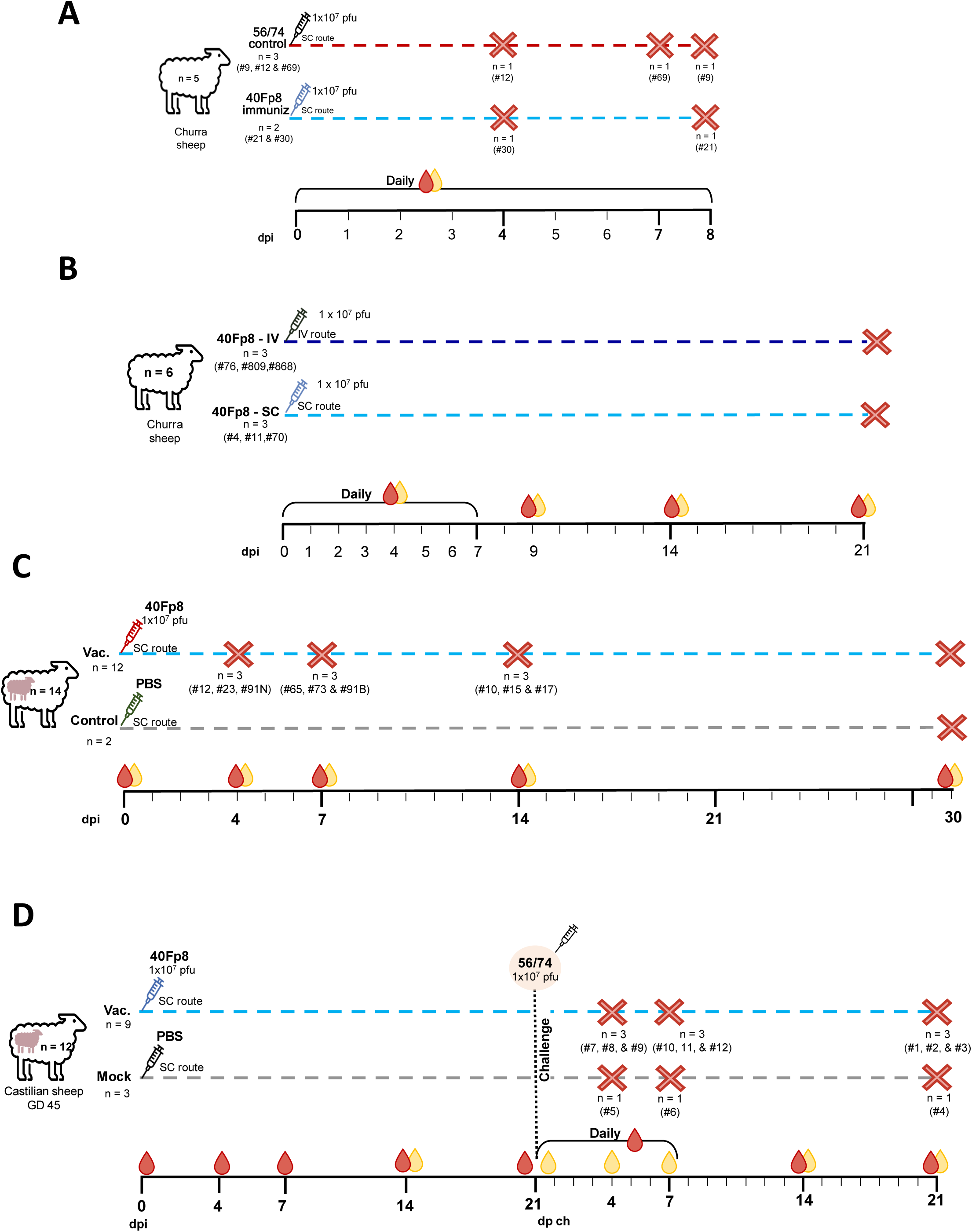
Experimental procedures in sheep for the analysis of immunogenicity (A & B), and safety (C) or protective efficacy (D) in pregnant animals of the vaccine strain 40Fp8. Sheep identification numbers are indicated. GD: gestational day. IV: intravenous inoculation. SC: subcutaneous inoculation. Dpi: days post-immunization. Dpc: days post challenge.

Four experiments were conducted to confirm the immunogenicity, safety and protective efficacy of the prototype live attenuated 40Fp8 vaccine against RVFV in non-pregnant and pregnant ewes of different Spanish breeds, as the sheep species is a natural host of the virus and highly susceptible to the disease *Experiment 1 (immunogenicity). Churra* ewes (n=5), aged 2 years, were used. In this experiment, two *Churra* ewes (#21 and #30) were inoculated subcutaneously (SC) in the neck area near the pre-scapular lymph node with 1 ml containing 10^7^ plaque forming units (pfu) of the RVFV-40Fp8 virus stock. These ewes were euthanized on days 4 and 8 post inoculation (dpi), respectively. A second group of three ewes (#9, #12 and #69) received identical dose of the parental RVFV strain 56/74. These animals were euthanized at 4, 7 and 8 dpi for pathological evaluation. Clinical evaluation was performed daily after inoculation. EDTA blood and serum samples were taken from the jugular vein (**figure 1A**).

*Experiment 2 (immunogenicity). Churra* ewe lambs (n=6), aged 4 months, were used. In this experiment, three animals (#4, #11 and #70) were SC inoculated in the neck area near the pre-scapular lymph node with 1 ml containing 1×10^7^ pfu of the 40Fp8 virus. Another three ewe lambs (#76, #809 and #868) were inoculated intravenously (IV) with an identical dose of 40Fp8. Animals were clinically monitored for three weeks after inoculation. EDTA blood and serum samples were taken daily during the first week or every other day (9, 11 and 13 dpi) during the second week. Two final blood and serum samples were taken at 14 and 21 dpi (**figure 1B**).

In experiments 1 and 2, the presence of virus in blood samples was examined by either virus isolation or RT-qPCR. Serum samples were used for biochemical analysis of enzymes indicative of hepatic injury and for detection of anti-nucleoprotein and/or anti-RVFV neutralizing antibodies. In experiment 2, antigen-specific cellular responses were also monitored by IFN-γ ELISA analysis after *in vitro* antigenic re-stimulation of EDTA blood samples taken at 14 and 21 dpi.

*Experiment 3 (vaccine safety).* Pregnant *Manchegan* ewes (n=14), between 3 and 5 years of age, oestrus-synchronized and naturally mated, were used. Pregnancy was confirmed by ultrasound prior to transferring the animals to the BSL-3 animal facility and by a pregnancy test on the day of inoculation (Idexx Rapid Visual Pregnancy Test kit). After the acclimatization period, ewes at gestational day 56 were inoculated SC with 1 ml containing 1×10^7^pfu of the RVFV-40Fp8 strain and euthanized in groups of n=3 on days 4 (#12, #23 and 91N), 7(#65, #73 and 91B), 14 (#10, #15 and #17) and 30 (#18, #26 and #40) post-immunization (dpi). Two pregnant ewes (#92 and #94) used as non-immunized controls were inoculated with mock-infected cell culture medium and euthanized at the end of the experiment (30 dpi). Throughout the experiment, clinical signs and rectal temperatures were examined daily. At the days of necropsy, blood and serum samples were collected for determination of viral loads (either by RT-qPCR or cell culture isolation) and for detection of anti-nucleoprotein and/or anti-RVFV neutralizing antibodies (**figure 1C**).

During the necropsies of the ewes and their foetuses, macroscopic evaluations of the lesions were performed. When present, lesions were scored according to their extent in the tissues and the severity of the lesion itself between 1 (focal/minimal lesions) and 4 (diffuse/severe lesions). To compare the level of foetal development, all foetuses found were measured and weighed. Samples were aseptically taken from the liver, spleen umbilical cord and placentomes of ewes, and from the liver, spleen and brain of foetuses. Amniotic fluid and blood from umbilical cords were also sampled. Additional samples were collected from any other organ in which macroscopic changes were observed. All samples were kept on ice during the necropsies and subsequently stored at −80 °C. Tissue samples were also fixed in 4% buffered formalin solution for 72 h, routinely processed and embedded in paraffin wax for subsequent routine histopathological studies (haematoxylin and eosin staining), as well as immunohistochemistry to visualize viral antigen using a polyclonal rabbit anti-RVFV serum, as described [19].

*Experiment 4 (vaccine protective efficacy).* Pregnant *Castilian* ewes (n=12), between 4 and 6 years of age, reproductively synchronized and naturally mated, were used. Pregnancy was confirmed by ultrasound prior to transferring the animals to the BSL-3 animal facility and by a pregnancy test on the day of inoculation (Idexx Rapid Visual Pregnancy Test kit). After the acclimatization period, nine ewes at gestational day 45 were inoculated with the RVFV-40Fp8 virus following the same immunization protocol as for experiment 3. Three pregnant ewes (#4, #5 and #6) were used as mock-immunized controls. Three weeks after immunization all animals were SC challenged with 1 mL containing 10^7^ pfu of the RVFV virulent strain 56/74. Throughout the experiment, animals were monitored clinically daily. Blood and serum samples were taken on different days after immunization (dpi) and after challenge (dpc) and used for the same assessments as described in experiment 3. Programmed necropsies were performed on 3 immunized ewes and 1 non-immunized control ewe at day 4 (ewe #5, #7, #8, #9), 7 (ewe #6, #10, #11, #12), and 21 (ewe #1, #2, #3, #4) post-challenge (dpc). The level of foetal development, macroscopic evaluations, tissue sampling and subsequent histopathological and immunohistochemical assessments were similar to those described for experiment 3.

### Blood chemistry

Blood samples were collected in BD Vacutainer tubes without anticoagulant. For separation of serum, the tubes were centrifuged at 1.267 × g for 10 min and stored at −80 °C until use. The sheep serum levels of aspartate aminotransferase (AST), gamma-glutamyltransferase (GGT) and lactate dehydrogenase (LDH), collected between 0 and 7 days post inoculation (experiment 1), were determined with a Saturno 100 analyzer, (Crony instruments) using specific reagents (BioAnalítica SL) according to the manufacturer’s instructions.

### Molecular, virological and serological assays

Different organ and blood samples were prepared and tested for the presence of RVFV RNA by RT-qPCR. Organ samples were processed to obtain 10% weight/volume homogenates in DMEM, by running one homogenization cycle (50 Hz oscillation frequency (50 cycles/s) for 4 minutes) in a Tissue lyser LT (Qiagen). Tubes were then centrifuged at 5.000 rpm for 5 min, supernatants collected and stored at-80°C until use. Viral RNA was isolated with the Speedtools system according to the manufacturer’s instructions (Biotools, Spain) from either 0.5 mL of blood or 0.5 mL of a 10% organ suspension. One to five microliters of purified RNA were subsequently used for RT-qPCR using the Scriptools RT-qPCR assay (Biotools, Madrid, Spain) in combination with an Eco-Real time PCR thermocycler (Illumina). One-tube RT-qPCR was performed using the forward L segment primer (5’-TTCTTTGCTTCTGATACCCTCTG-3’), L segment reverse primer (5’-GTTCCACTTCCTTGCATCATCTG-3’) and a FAM-BHQ1-labelled probe (5’-FAM-TTGCACAAGTCCACACAGGCCCCT-BHQ1-3’) as described [20].

*In vitro* virus isolation was attempted in those RT-qPCR positive samples to confirm or to rule out the presence of infectious virus. Briefly, 0.1-0.5 mL of whole blood or clarified organ homogenates was adsorbed for 1 hour to semi confluent Vero cell monolayers in 25 cm^2^ tissue culture flasks. After adsorption, cells were extensively washed with PBS and incubated for 5 days. In case of absence of cpe, this procedure was repeated up to three blind passages.

Detection of RVFV serum nucleoprotein N-specific antibodies was determined by an IDVet RVF screen Kit (IDVet, France). The titers of RVFV serum neutralizing antibodies (nAbs) were determined using a virus microneutralization test (VNT). Briefly, in a 96-well plate format, serial dilutions (50 μl) of heat-inactivated sera (2 h, 56 °C), in 2-4 replicas, were incubated with 50 μl of the attenuated RVFV-56/74 or 40Fp8 strain (100 TCID50/ml) for 2 h at room temperature. Subsequently, 2,5×10^4^ Vero cells (in 50 μl) were added to each well. Plates were incubated for 3-4 days at 37 °C and 5% CO2 and cytopathic effect (CPE) scored under microscope observation. VNT50 titres were calculated using the Reed and Muench 50% end-point method.

### IFN-γ detection assay

An *in vitro* re-stimulation assay was performed in MW24 plate wells using 0.5 mL of whole EDTA blood samples per well, which were incubated with 1 µg/mL of either recombinant purified proteins Gn, or N, or 5 µg of Gn-derived peptides #19 (203-FQSYAHHRTLLEAVH-217) and #21 (211-TLLEAVHDTIIAKAD-225), or Gc-derived peptides #226 (1031-AAFLNLTGCYSCNAG-1045) and #253 (1139-SWNFFDWFSGLMSWF-1153) [18]. The plates were incubated at 37 °C in a 5% CO2 incubator for three days. The plasma was recovered by centrifugation of plates (300 x g) and tested in a IFNγ capture sandwich ELISA assay as described [18, 19]. Briefly the plasma from both SC and IV inoculated ewes (experiment 2) were collected at 14 and 21 dpi and added to the ELISA plate wells previously coated with anti-IFNγ mAb purified (MT17.1, Mabtech). Upon incubation and washing steps a detector anti-bovine IFNγ mAb conjugated with biotin (MT307, Mabtech) was added. Streptavidin-HRPO (Becton-Dickinson) was used for detection of immunocomplexes upon addition of TMB peroxidase substrate (Sigma-Aldrich). Absorbance values were determined at 450 nm by an automated plate reader (BMG, Labtech).

### Data Visualization and Analysis

This study was exploratory in nature, using a low number of individuals for complying with ethics, welfare and strict biosafety regulations. Statistical methods were not applied due to the insufficient sample size considering high variability between subjects. We focused instead on describing trends and patterns rather than seeking statistical significance. The valuable information obtained in this work is therefore based mainly on descriptive and qualitative analyses. The graphical representation of the results, both individual and mean values, was carried out with GraphPad Prism Version 7.0 (GraphPad Software, La Jolla, CA, USA)

## Results

### Preliminary evaluation of the safety and immunogenicity of a high dose of 40Fp8 strain inoculation in ewes (experiment 1)

We previously demonstrated that the 40Fp8 virus was highly attenuated in mice. To confirm attenuation in a natural host, a high dose of 40Fp8 (10^7^ pfu) was inoculated subcutaneously in ewes (**figure 1A**). Then, clinical, virological, immunological and biochemical parameters were compared with control ewes that received the same dose of the parental virulent RVFV strain 56/74. There was a clear increase in rectal temperature at 1 dpi that peaked at 2 dpi in ewes that received 56/74 virus (#9, #12, #69). In contrast, neither of the two ewes inoculated with 40Fp8 (#21, #30) showed hyperthermia at any time after inoculation (**figure 2A**). Isolation of infectious virus was achieved in two of the three ewes on the first day after inoculation with 56/74, and in all three ewes at 2 and 3 dpi, while no infectious virus was detected in ewes inoculated with 40Fp8 throughout the experiment (**figure 2B**). AST, GGT and LDH serum levels, indicative of liver/tissue damage, increased from 2 dpi in ewes inoculated with 56/74 in contrast to animals receiving 40Fp8 (**figure 2C)**. In the group inoculated with 40Fp8 it was possible to detect induction of specific antibodies although slightly delayed and of lower magnitude with respect to the group inoculated with 56/74 (**figure 2D**). Taken together, the absence of clinical disease, viremia and hepatic damage markers in ewes inoculated with a high dose of 40Fp8 virus, together with the specific and neutralizing antibody responses, confirmed its attenuated phenotype and suggested its immunogenic character.

**Fig. 2.**
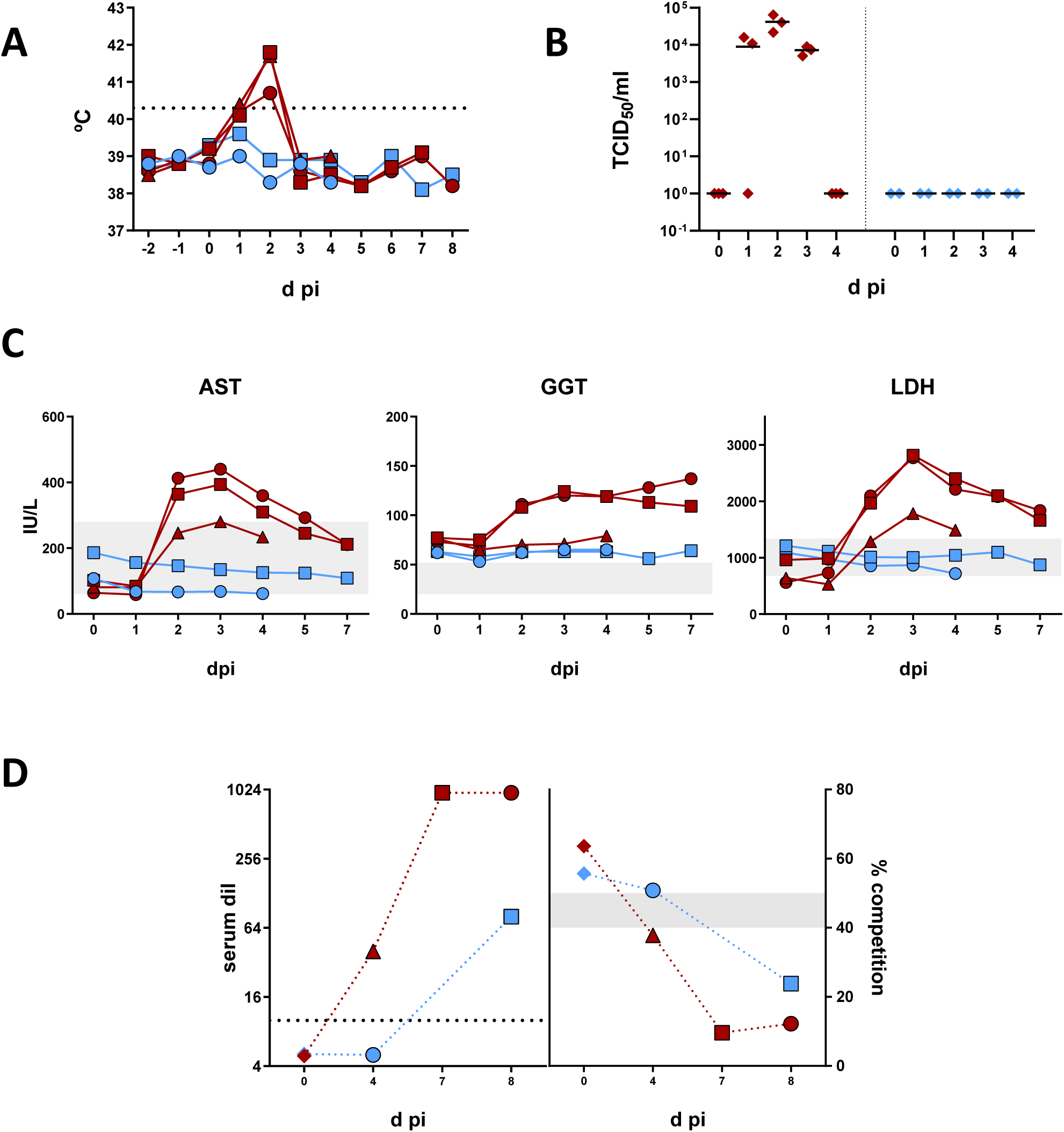
Comparative evaluation of rectal temperatures, viremia levels, liver enzymes and RVFV-specific antibodies in ewes inoculated with 40Fp8 or 56/74 strain. **A.** Rectal ewes temperature records before and after virus inoculation. **B.** Viremia levels obtained by titration in cell culture. The values displayed correspond to 50% tissue culture infectious dose per mL (TCID50/mL). The horizontal black dotted line marks hyperthermia. **C.** Kinetics of hepatic enzymes plasmatic levels after inoculation of each sheep. The shaded areas depict normal range values. Values given in International Units (IU) per mL. **D.** Kinetics of serum antibody responses. Each symbol represents values of individual sheep at the days of sacrifice (4, 7, 8). Left panel: neutralizing antibodies. The values correspond to the reciprocal of serum dilution causing a 50% reduction of infectivity in cell culture. Right panel: anti-nucleoprotein antibodies. Values correspond to competition percentages as determined by the IdVet ELISA test. Shaded area between 40-50% indicates the doubtful range. Red symbols: sheep inoculated with the wild-type 56/74 strain. Blue symbols: sheep inoculated with 40Fp8 virus.

### Comparative evaluation of the immunogenicity induced by a high dose of 40Fp8 strain administered by different routes of inoculation in ewes (experiment 2)

An additional 40Fp8 inoculation experiment was performed in three ewes (#76, #809, #868) using the intravenous (IV) route instead, with a high dose of 1.0 x 10^7^ pfu (3,2 x10^7^ after back titration). In parallel, three ewes (#4, #11, #70) were also SC inoculated with the same dose to compare the effect of the route of administration (**figure 1B**). On the same day of inoculation (d0), the sheep #11, included in the group to be inoculated subcutaneously, showed a slight hyperthermia. However, no further temperature increases were observed after inoculation with 40Fp8 in any of the animals (**figure 3A**). In general, the values of biochemistry parameters (hepatic transaminases, total protein and blood urea nitrogen) showed no significant alterations with respect to normal levels, as observed in the previous experiment and in contrast to the values usually found in ewes inoculated with the virulent RVFV 56/74 isolate [18]. Assessment of RNAemia by RT-qPCR rendered negative Cq values (≥40). Only one blood sample (day 5, ewe #809, IV route) rendered a Cq value below 40 (35.88). However, viral isolation attempts using blood from this sheep were negative upon three consecutive blinded passages in cell culture, indicating a very low or null systemic dissemination of 40Fp8 irrespective of the delivery route. By days 5-7 post inoculation, most of the ewes’ sera had detectable nAb titers at or above the threshold values of the assay. These data also suggested active virus replication, as confirmed by the kinetics of anti-N antibody induction by competition ELISA (**figure 3B**), also confirming the proper induction of virus-specific antibody responses. No significant differences were found in the kinetics of induction of humoral immunity in relation to the route of administration. To assess virus-induced cellular immunity, an IFNγ ELISA assay was performed after re-stimulation of whole blood collected at 14 or 21 dpi. IFNγ secretion was clearly detected after re-stimulation with Gn and Gc specific peptides in 2 of the ewes inoculated IV but only at 21 dpi, and in both groups when recombinant proteins (Gn or N) were used as stimuli (**figure 3C**). Taken together, these data point to the ability of 40Fp8 strain to elicit a complete immune response in ewes, despite its high level of attenuation.

**Fig 3.**
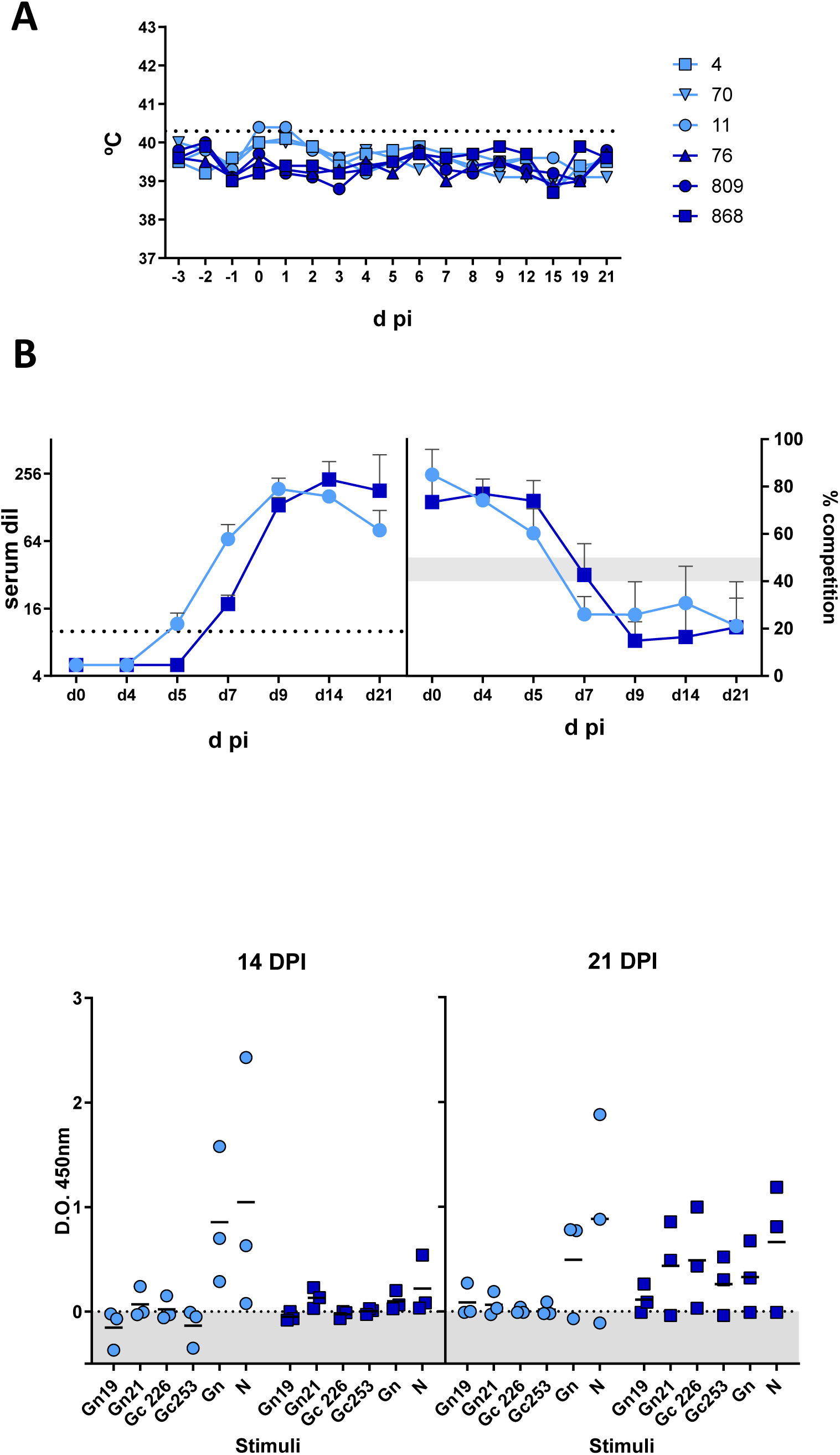
**A.** Evolution of rectal temperature in ewes after SC (light blue) or IV (dark blue) inoculation with 40Fp8 virus. The horizontal black dotted line marks hyperthermia. **B.** Serological determination of neutralizing antibody titers by TCID50 microneutralization assay (left panel) and competition ELISA anti-N (right panel). Both graphs show mean plus SD values. The dotted line depicts the sensitivity threshold (1/10 dilution). The grey shaded area depicts the doubtful range. **C.** Detection of IFNγ in plasma by capture ELISA. Blood samples taken at 14 or 21 days after SC or IV 40Fp8 inoculation were re-stimulated with recombinant RVFV Gn or N proteins or peptides derived from the Gn (#19 and #21) or Gc (#226 and #253) protein sequences. Each symbol denotes individual values (after subtraction of values derived from unstimulated plasma) and horizontal bars denote mean values.

### Vaccine safety study in pregnant ewes (experiment 3)

Since inoculation of 40Fp8 was immunogenic both at the humoral and cellular level in ewes, we were then aimed to test its vaccine potential in this host species using a more relevant and susceptible animal model such as pregnant sheep. RVF causes stillbirths and abortions in pregnant ruminants, therefore a live attenuated vaccine should be free of any residual virulence that could induce pregnancy failure in such susceptible hosts. To rule out residual virulence, we administered a subcutaneous high dose of 40Fp8 in 12 pregnant ewes at gestational day 56 when sheep are more susceptible to reproductive failure (**figure 1C**). Apart from a mild temperature increase on day 1 after immunization **(figure 4A)**, the ewes showed no clinical signs or reproductive failure leading to foetal death, congenital malformations, resorptions or miscarriages during the experiment. During necropsies carried out between 4 and 30 dpi, the ewes showed no relevant macroscopic or histopathological lesions in target organs of the virus (placentomes and liver) (**supplementary figures 1 & 2**) and the developmental parameters of the foetuses were as expected for their gestation period (**figure 4B**). No significant differences were observed in weight or length of foetuses obtained from immunised ewes and control ewes at the end of the experiment (30 dpi; 86 days of gestation). In addition, all foetuses obtained throughout the experiment from immunised ewes as well as those from control ewes at the end of the experiment, presented a normal appearance during macroscopic evaluations (**figure 4C**).

**Fig. 4.**
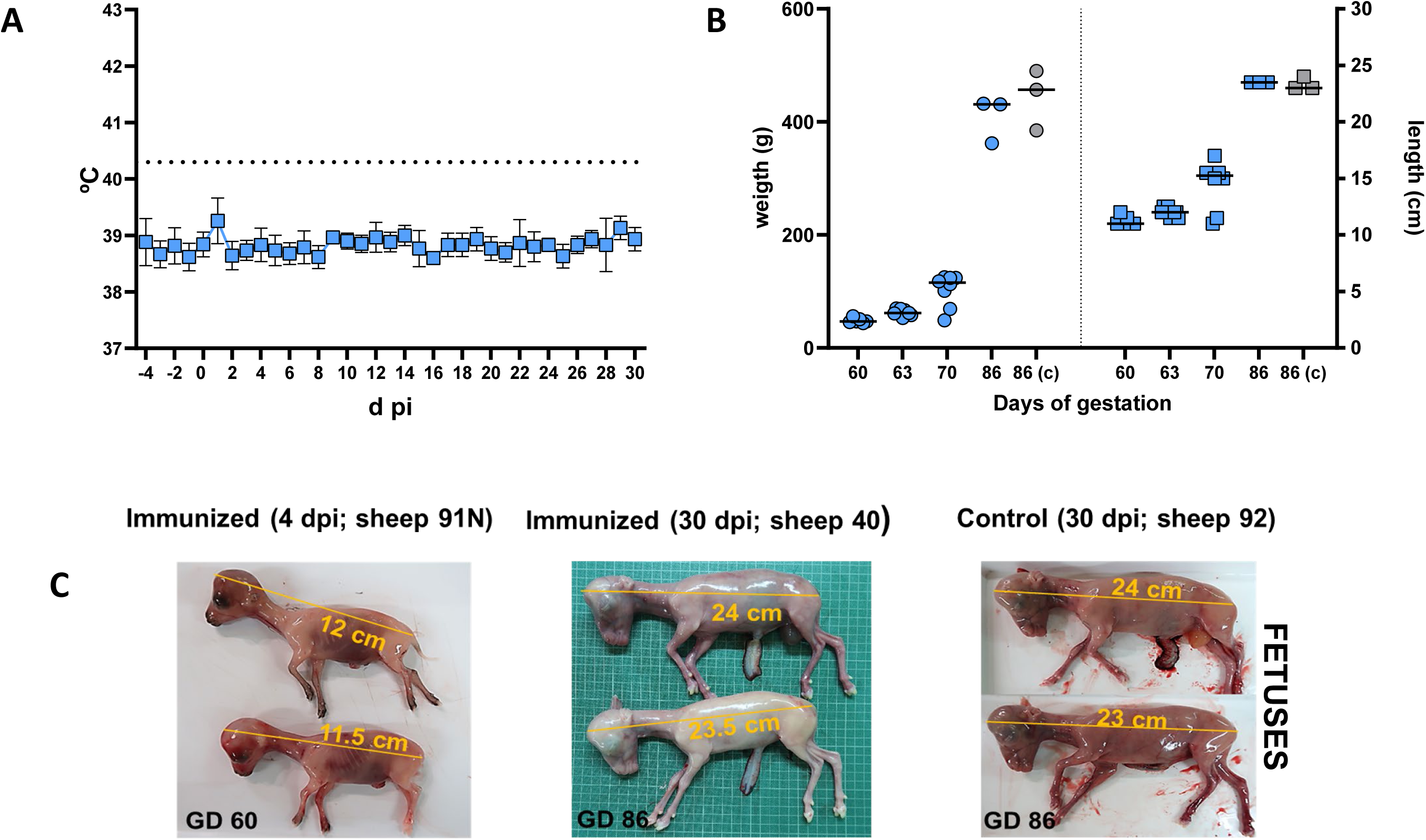
**A.** Rectal temperature kinetics (mean ± SD) in pregnant ewes inoculated subcutaneously with 10^7^ pfu of the 40Fp8 virus. The horizontal dotted line marks hyperthermia. **B.** Evaluation of foetal development. Mean and individual values for size and weight of foetuses found after necropsy at different days of gestation (GD). **C.** Representative images of foetuses at different days post inoculation (dpi) in immunized or control ewes.

Only low levels of viral RNA were detected by qPCR in some animals in the early stages after immunization in blood samples from umbilical and jugular veins (ewes) (**figure 5A)**. The viral genome was practically undetectable from 14 dpi onwards. The presence of the viral genome was also assessed in tissue samples obtained from foetuses (liver, spleen and brain) and ewes (liver, spleen and placentomes) where levels were also very low or undetectable throughout the experiment (**figure 5B**). Accordingly, no infectious virus was isolated from any of these samples with Cqs over the threshold after three blind serial passages in Vero cell cultures. Additional histopathological studies revealed no remarkable histopathological lesions in any of the tissues evaluated, including target organs of the virus (liver and placentomes). Immunohistochemical evaluations also did not reveal the presence of viral antigen in the tissues examined.

**Fig. 5.**
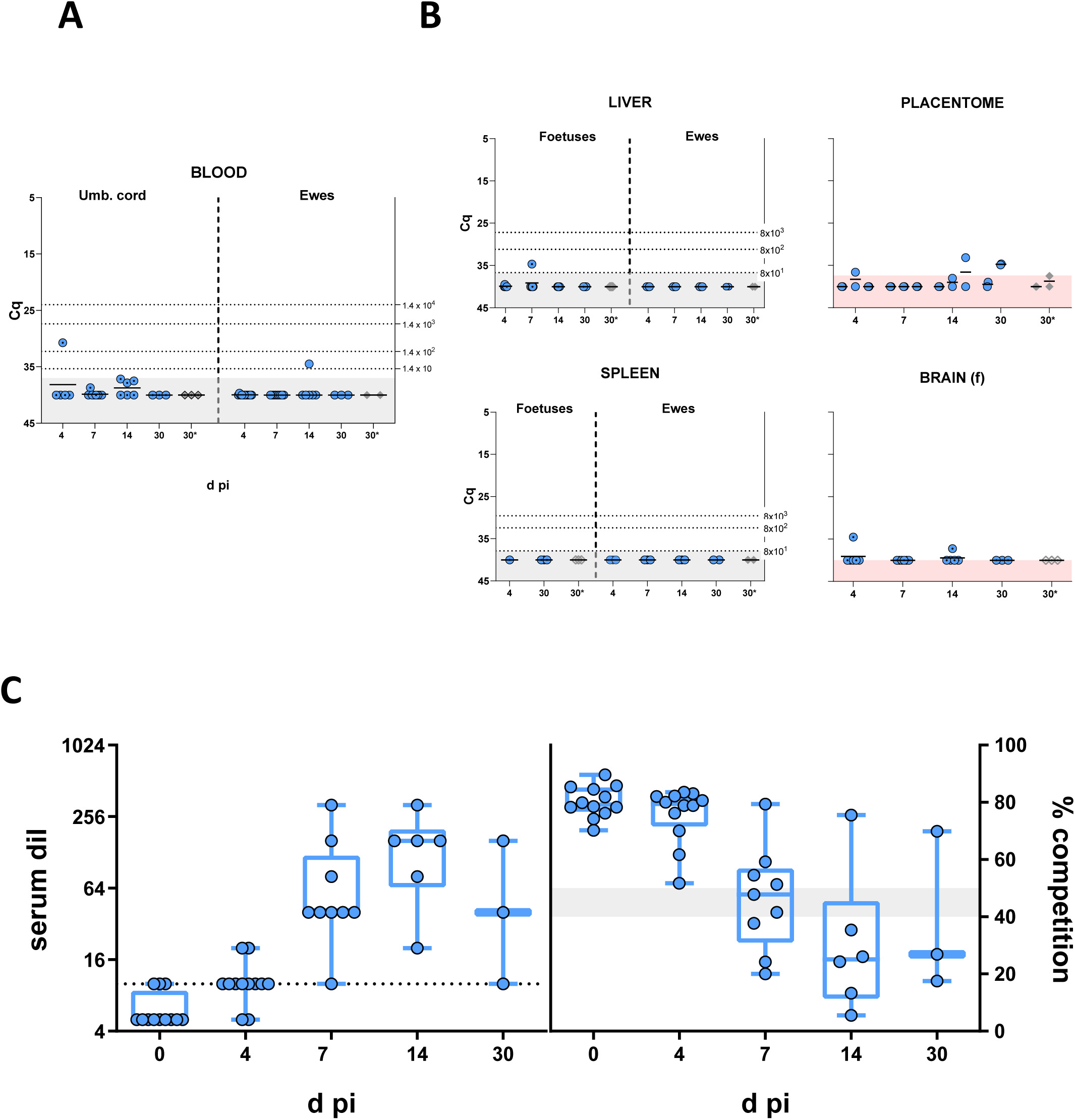
**A.** Detection of viral RNA by RT-qPCR in blood samples taken from umbilical (foetus) or jugular veins of pregnant ewes at various days after inoculation with the 40Fp8 virus. **B.** Detection of viral RNA by RT-qPCR in foetal and/or maternal tissues at various times after inoculation with the 40Fp8 virus. Data from liver, spleen and placentome (cotyledon) collected from pregnant ewes as well as data from liver, spleen and brain taken from foetuses are shown. The shaded areas indicate the sensitivity limit of the assay. The dotted lines indicate the ratio between the Cq obtained and the corresponding viral infectious load or pfu equivalents. **C left.** Kinetics of neutralizing antibody levels assessed by microneutralization plate in sheep sera obtained on the indicated dates after vaccination with 40Fp8 virus. The dotted line indicates the sensitivity limit of the assay. **C right.** Kinetics of induction of anti-nucleoprotein N antibodies assessed by competitive ELISA (IDVet RVFV screen test). Median and interquartile range as well as individual values are represented. The shaded area indicates the range of doubtful samples.

Immunised ewes seroconverted between 4 and 7 dpi, as shown by the kinetics on anti-nucleoprotein antibodies in serum (**figure 5C**), developing a clear neutralising antibody response from 4 dpi, which reached levels (serum dilution over 1/40) correlating with protection from 7 dpi onwards. These results demonstrated that a high dose of RVFV-40Fp8 did not cause clinical signs or reproductive failure in pregnant ewes, but induced levels of immunity correlated with protection, thus appearing as a safe vaccination approach against RVFV.

### 40Fp8 vaccine protective efficacy study in pregnant ewes (experiment 4)

The previous data advocated for the use of 40Fp8 strain as a promising vaccine candidate since pregnant ewes immunized with 40Fp8 were free of adverse effects and developed nAbs from days 4-7. To test whether the 40Fp8 strain induced immunity was indeed sufficient to protect both ewes and foetuses from infection, a new group of pregnant ewes at gestational day 45 was inoculated with 10^7^ pfu of 40Fp8, and three weeks after vaccination were challenged subcutaneously with 5×10^6^ pfu of the virulent RVFV 56/74 strain (**see experimental outline fig 1D**). To evaluate the occurrence of lesions that could affect both pregnant ewes and foetuses after challenge, groups of vaccinated and non-vaccinated control animals were necropsied sequentially on days 4, 7 and 21 post-challenge (dpc). When present, lesions were scored according to their extent in the tissues and the severity of the lesion itself between 1 (focal/minimal lesions) and 4 (diffuse/severe lesions). One vaccinated ewe (#9) scheduled to be euthanized on day 4 post challenge developed hyperthermia on day 21 post immunization (**figure 6A**) and aborted prior to challenge. To ascertain if this abortion was induced by 40Fp8 vaccination, the aborted foetus and placental residues recovered from this ewe were subjected to RT-qPCR as well as isolation of RVFV in cell culture, rendering negative results in both assays (not shown). To further investigate the possible cause of this outcome, the aborted tissue samples were also analyzed by qPCR for the presence of several abortifacient pathogens (i.e., Border disease virus, *Chlamydia abortus, Coxiella burnetii, Toxoplasma gondii, Neospora caninum, Listeria monocytogenes, Campylobacter fetus, and Brucella spp*). No positive hit was found to identify a causative pathogen for this outcome. On day 2 after challenge with the virulent RVFV strain 56/74, all mock-immunized ewes (#4, #5 and #6) displayed higher temperatures than immunized ewes, with values above 40°C for ewe # 6. Ewe #5 also exhibited temperatures above 40°C on 4 dpc (**figure 6A**). At necropsy, following a macroscopic pathological analysis, non-immunised control ewes euthanized at 4 (ewe #5) and 7 dpc (ewe #6) exhibited severe scattered foci of necrosis in the liver and increased volume of amniotic fluid (**figure 6B-C**). Although the growth parameters of their foetuses were normal, the foetuses of control ewe #6 euthanized at 7 dpc were found dead and were collected in imminent abortion (**figure 6C**). Non-immunized control ewe #4 aborted on 17 dpc, being euthanized that same day. At necropsy, there was evidence of vaginal mucopurulent discharge associated with acute necrotic haemorrhagic endometritis, without the presence of macroscopic hepatic lesions. Remains of placenta, but not of the foetuses, were found. In contrast, no liver or reproductive organ lesions were observed in any of the immunized ewes euthanized throughout the experiment, and their foetuses showed normal growth parameters and appearance (**figure 6C**).

**Fig. 6.**
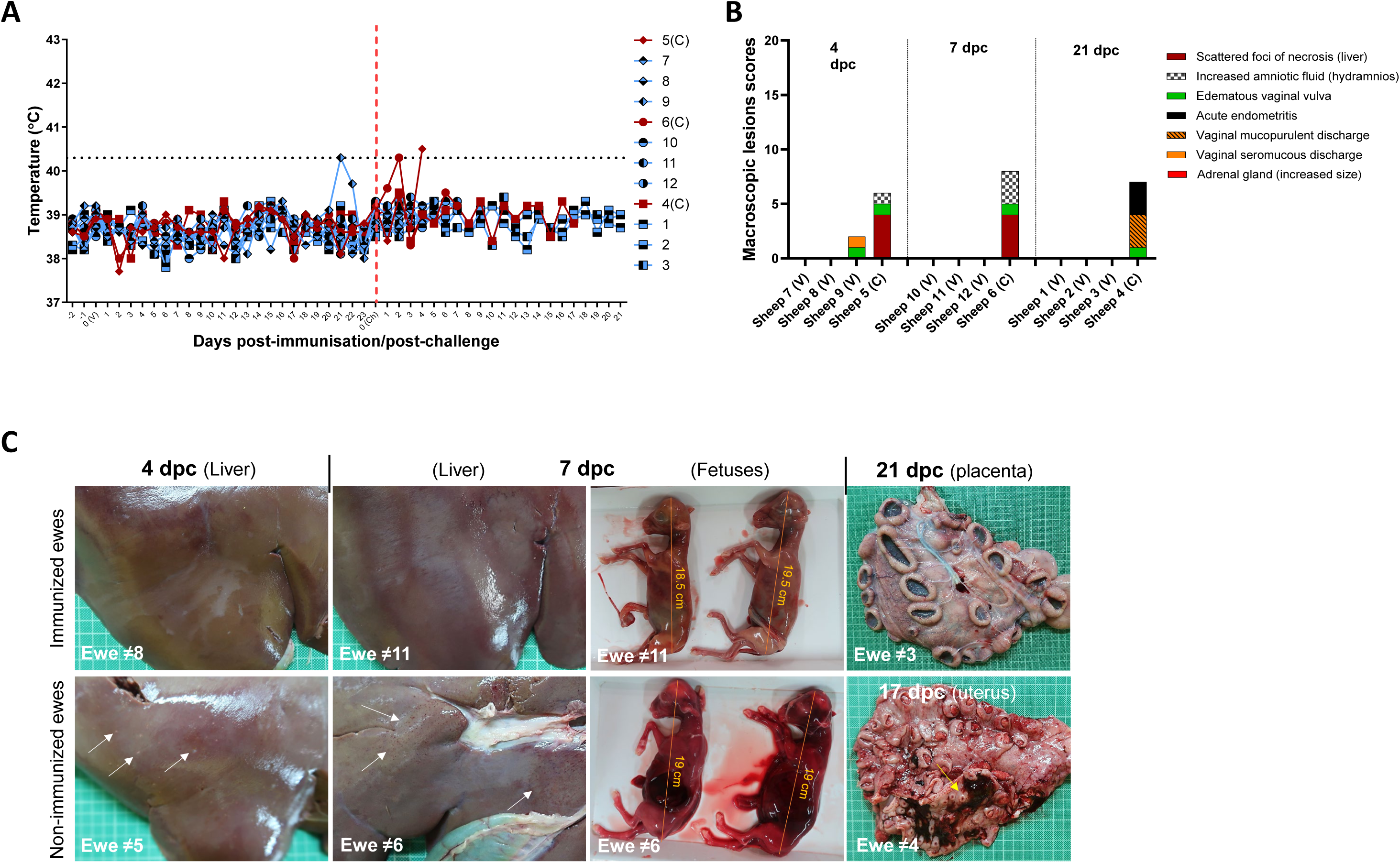
**A.** Rectal temperatures of individual ewes before and after virus challenge (red vertical dotted line). The horizontal black dotted line marks hyperthermia. **B.** Macroscopic lesions and scores observed in sheep upon 56/74 challenge. When present, lesions were scored according to their extent in the tissues and the severity of the lesion itself between 1 (focal/minimal lesions) and 4 (diffuse/severe lesions). **C.** Representative images of liver, foetuses and placentas found in immunized (upper images) and non-immunized sheep (bottom) after challenge. Scattered foci of necrotic lesions were found in the livers of non-immunized control ewes (white arrows) along with the presence of foetuses found dead and collected in imminent abortion. In contrast, livers without lesions and live foetuses were found in the 40Fp8 immunized ewes. Acute purulent metritis (yellow arrow) was observed in a non-immunized control ewe that aborted at 17 dpc, in contrast to the normal aspect placenta of a protected immunized ewe that was euthanized at 21 dpc.

Histopathological evaluation of the livers of non-immunized ewes euthanized at 4 (ewe #5) and 7 dpc (ewe #6) revealed findings characteristic of RVFV infection, with multifocal necrotic foci filled with cellular debris and both degenerated and viable neutrophils. Viral infection was also confirmed via immunohistochemistry, with hepatocytes within the necrotic foci appearing as the main cells immunolabelled against viral antigen (**figure 7A,E**). Viral antigen was also occasionally observed in the placentomes at 4 dpc (non-immunized ewe #5), specifically in maternal epithelial cells (**figure 7B-D**). However, at 7 dpc, antigen-positive cells were visible throughout the placentomes of non-immunized ewe #6, both in the maternal epithelial cells/syncytial cells and in the foetal trophoblasts (**figure 7F,G**). These cells, mainly maternal epithelial cells, were enlarged and showed necrosis accompanied by viable, degenerated neutrophil infiltrates. Neutrophil infiltrates were also often observed at the base of the maternal villi. As for the tissues collected from non-immunized ewe foetuses at 4 and 7 dpc, viral antigen was only detected in brain, kidney and liver of foetuses in imminent abortion sampled at 7 dpc from ewe #6 ewe (**figure 7H-M**).

**Fig 7.**
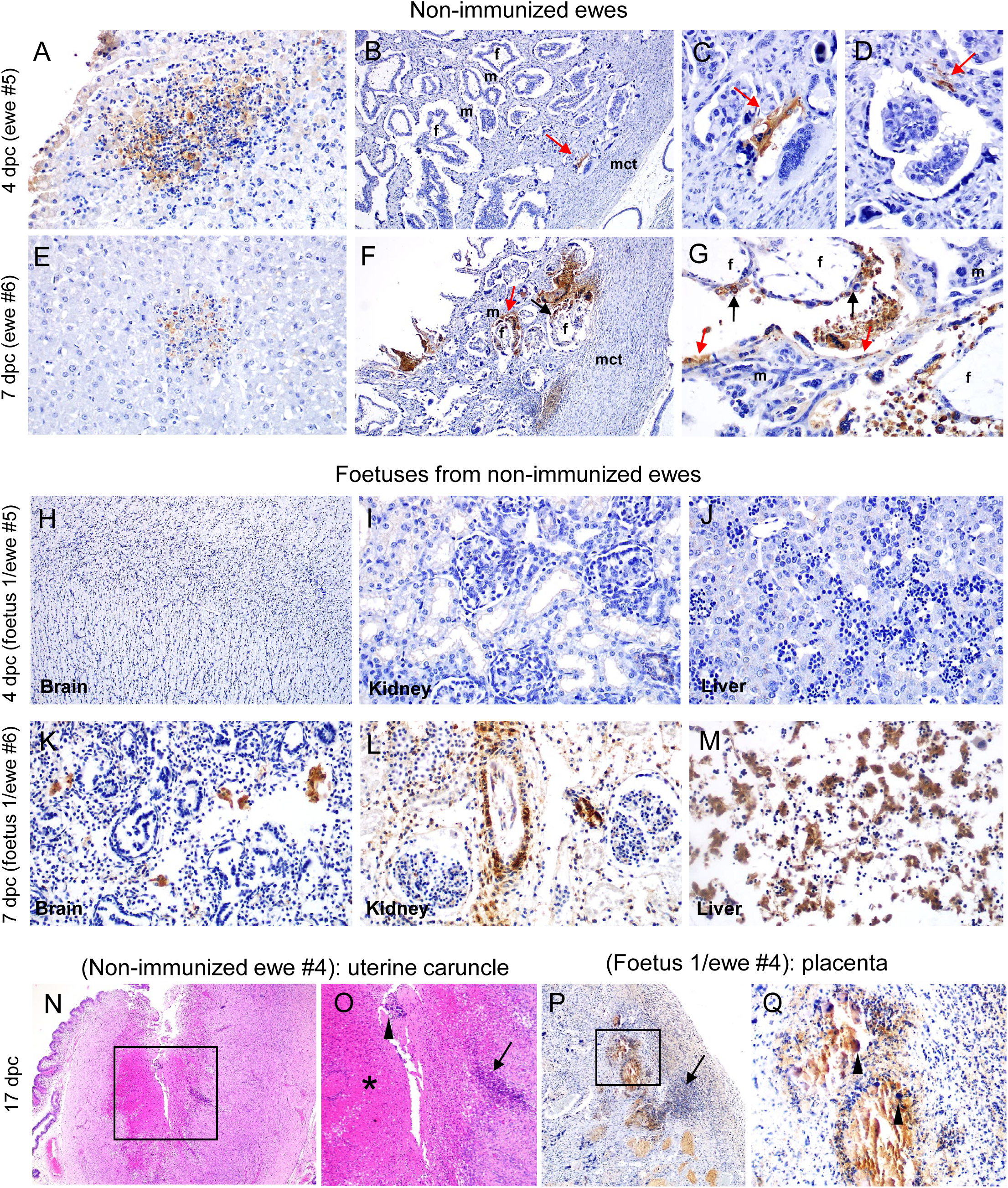
Representative images of histopathological lesions (Haematoxylin-eosin staining) and immunohistochemical detection of RVFV antigen in different tissues obtained from pregnant ewes not immunized with the 40Fp8 strain and their fetuses at different days after challenge. Note the presence of hepatocytes immunolabelled against viral antigen within necrotic foci in the liver of non-immunized ewes at 4 and 7 dpc **(A,E, 40x).** Viral antigen was also occasionally observed in the placentomes at 4 dpc **(B, 10x; C,D, x40x),** specifically in maternal epithelial cells (red arrows). However, at 7 dpc, antigen-positive cells were visible throughout the placentomes of non-immunized ewe #6, both in the maternal epithelial cells/syncytial cells (red arrows) and in the foetal trophoblasts (black arrows) **(F, 10x; G, 40x).** Viral antigen was detected in brain, kidney and liver of foetuses in imminent abortion sampled at 7 dpc from ewe #6 ewe, but not in foetuses sampled at 4 dpc **(H-M, 40x).** Caruncular endometrium of ewe #4 aborted on 17 dpc revealed the presence of areas of coagulative necrosis and haemorrhages (asterisk), foci of mineralization (black arrowhead) and lymphoplasmacytic and neutrophilic infiltrates (black arrow) **(N, 10x; O, magnified area, 40x).** Remains of placenta from the aborted foetus of ewe #4 **(P, 10x; Q, magnified area, 40x).** Note the presence of lymphoplasmacytic and neutrophilic infiltrates (black arrow) as well as the presence of viral antigen in the placental epithelial cells (black arrowheads). **(f)** foetal villus; **(m)**; maternal villus; **(mct)**: maternal connective tissue.

**Figure 8.**
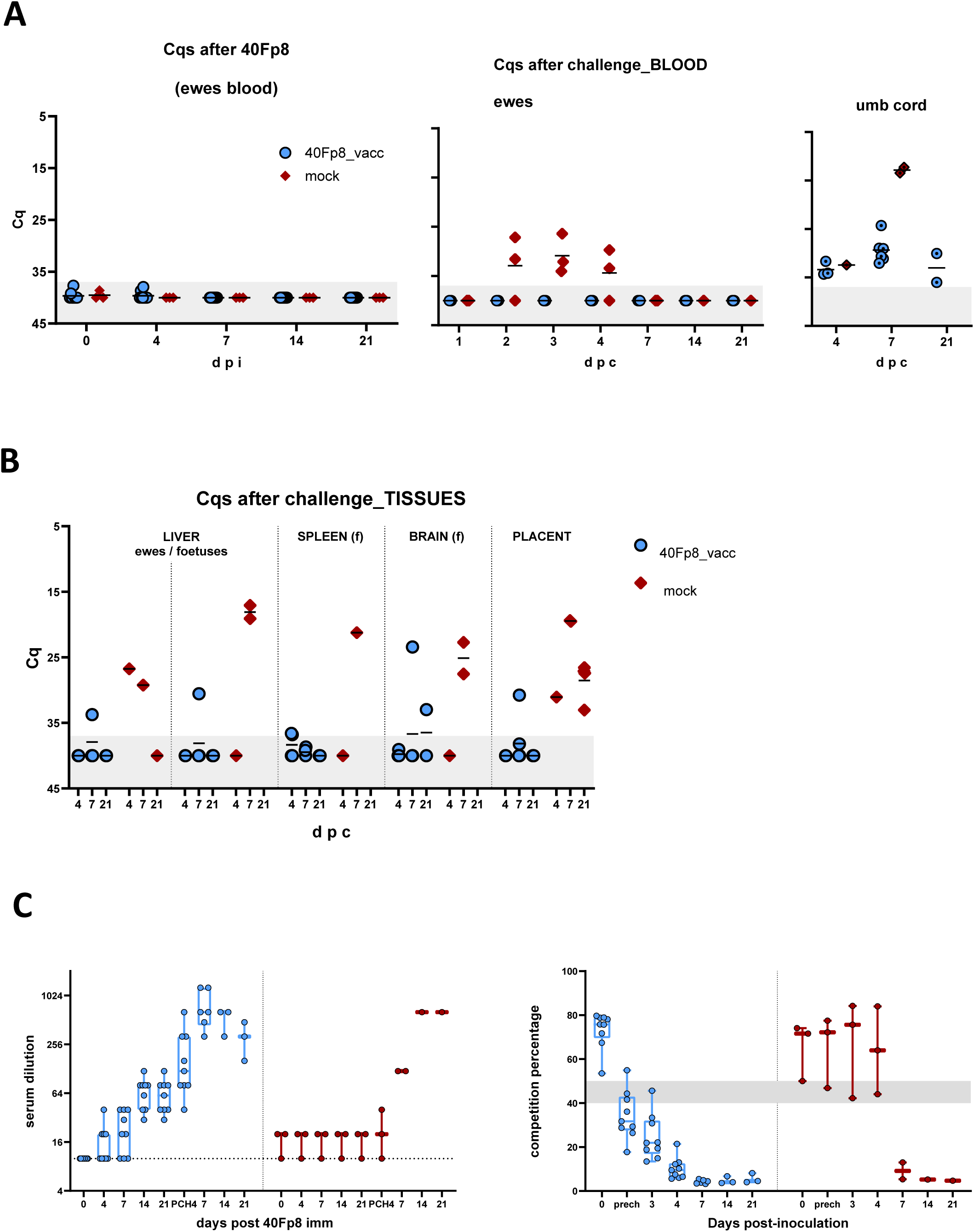
**A.** RT-qPCR analysis of viral RNA in vaccinated ewes and foetal (umbilical vein) blood before and after challenge. **B.** Detection of viral RNA in sheep (liver and placentome) or foetal (liver, spleen, brain) tissue samples by RT-qPCR. In A and B, the shaded area indicates the sensitivity limit of the assay. **C** (**left)**. Neutralizing anti-RVFV antibodies (nAbs) detected in sera by virus microneutralization assay in cell culture. The dotted line indicates sensitivity threshold. **C (right).** Detection of serum anti-nucleoprotein antibodies by competitive ELISA (IDVet RVF screen Kit). The shaded area indicates the estimated range of doubtful samples. In C and D median and interquartile range (for n >3) as well as individual values are represented.

Histopathological evaluation of the caruncular endometrium of ewe#4 aborted on 17dpc revealed the presence of áreas of coagulative necrosis with loss of tissue architecture and haemorrhages, foci of mineralization, focally extensive to multifocal lymphoplasmacytic and neutrophilic infiltrates together with fibrosis at the periphery of the lesion (**figure 7N,O**). The placental remains found corresponding to the aborted foetus of ewe #4 also showed areas of necrosis together with diffuse lymphoplasmacytic and neutrophilic infiltrates. By immunohistochemistry, viral antigen was detected in the placental epithelial cells of this aborted foetus, but not in the uterine caruncles of ewe #4 (**figure 7P,Q**). Finally, the liver of ewe #4 showed neither lesions nor presence of viral antigen.

In contrast, tissue samples taken from 40Fp8 vaccinated ewes and their foetuses showed no remarkable histopathological lesions and no presence of viral antigen, which evidences that vaccination with 40Fp8 controls the spread of RVFV to reproductive tissues and foetuses.

Analysis of blood samples for RVFV RNA detection was performed by RT-qPCR. As expected, viral genome was not detected in any of the blood samples taken after immunization with 40Fp8. After challenge, all immunized ewes remained negative, and only samples from non-vaccinated ewes yielded positive results in the 25-35 Cq range between 2-4 dpc (**figure 9A**). These RNA-positive blood samples yielded positive virus isolation in Vero cell culture after the first passage, while samples from immunized ewes were negative for virus isolation after three passages. Blood collected from umbilical vein during necropsies was also subjected to viral RNA detection. Cq values above the limit of detection were found in all foetuses, but upon virus isolation no infectious virus was recovered except for mock immunized ewes’ foetal samples at 7 dpc **(figure 9A**). Liver tissue was also evaluated by RT-qPCR for the presence of RVFV RNA in both ewes and foetuses. The lowest Cq values corresponded to the livers of control ewes tested in the 4 and 7 dpc groups. Livers from foetuses carried by RVFV-RNA positive ewes were also RNA-positive on day 7. Liver samples from vaccinated ewes and their foetuses displaying Cq values above threshold levels were negative to viral isolation attempts. Other tissues obtained from vaccinated animals (placentome, foetal brain and spleen) were evaluated and in all cases yielded negative to virus isolation, despite some background RNA detection by RT-qPCR. In contrast, viral isolation was successful in those samples tested on foetuses carried by non-immunized control ewes (**figure 9B**). These results agree with viral antigen detection described by immunohistochemistry.

As expected, by day 7-14 all vaccinated ewes had seroconverted with protective nAbs titers that increased upon the virulent challenge (**figure 9C**). The kinetics of anti-N antibodies followed a similar pattern, indicating effective priming of the immune system by the vaccine candidate (**figure 9D**)

## Discussion

One of the hallmarks of RVF outbreaks in Africa is the massive occurrence of abortions in sheep flocks [21]. During the breeding seasons nearly 100% of the pregnant sheep abort upon a RVFV infection, resulting in heavy economic losses and social consequences [22–24]. While vaccination can contribute to contain the impact of the disease, vaccines should be safe to foetuses and capable of provide enough level of maternal protection after immunization to avoid the spread of the virus into foetal and reproductive tissues. If vaccination is carried out using a live attenuated vaccine (LAV), it should not retain residual virulence to ensure the completion of the gestational cycle. Several LAVs such as the Smithburn, clone 13 or MP12 can be safely administered in adult sheep but their use during pregnancy has been questioned due to potential adverse effects on the foetuses [25–27].

LAVs can be optimal candidates for livestock vaccination programmes; however, the potential for reassortment events, reversion to (or the presence of residual) virulence has frequently been identified as concerns that could potentially contribute to disease spread, thereby limiting their use in immunosuppressed hosts or in pregnant animals [28–30]. To overcome these problems, novel, rationally designed, reverse genetics-based vaccine candidates as well as other strategies have been recently developed and some of them are in early phases of clinical developments [31].

We showed in previous works that the mutagenized RVFV variant 40Fp8 virus is highly attenuated even in an immunodeficient mouse model [12, 13]. We showed here that the inoculation with the 40Fp8 virus does not alter the basal biochemical parameters, in sharp contrast to what occurs when a virulent strain is inoculated through the same route in adult sheep, indicating that 40Fp8 can be administrated safely in sheep. However, a high level of attenuation could hamper the elicitation of an effective immune response. Therefore, here we tested firstly the immunity induced in sheep to assess its potential as a novel candidate vaccine. This was performed in two pilot experiments using a reduced number of sheep. Taken together, it was possible to confirm that at the doses used here, the parenteral inoculation of 40Fp8 was able to trigger an immune response of a sufficient magnitude to induce a neutralizing antibody response to titres usually over the described protective threshold. This is important since the induction of neutralizing antibodies is a clear correlate of protection against RVF [32–34]. However, the minimal immunization dosage of 40Fp8 able to induce protective antibody titers in the sheep model should be further investigated. In previous works, RVF LAV doses of 10^4^-10^5^pfu were shown to be protective in gestational sheep models [35, 36]. In our hands, doses of 40Fp8 ranging 10^3^-10^4^ pfu, fully protected mice from a virulent challenge (unpublished observations). Due to the highly attenuated character of 40Fp8 it may be necessary to inoculate with high virus titers to ensure full protection using a single dose only. Of note, our results also show that the inoculation of high doses of the attenuated 40Fp8 virus induces cell-mediated immune responses since IFN-γ was detected after *in vitro* re-stimulation of 40Fp8-immune blood cells. Nonetheless, a more in-depth characterization will be needed, to ascertain if memory T-cell responses can be also detected long after immunization.

The subcutaneous immunization of pregnant ewes with RVFV-40Fp8 induced immunity levels which correlated with protection in the absence of clinical disease and low to undetectable levels of viral RNA. After virulent RVFV-56/74 challenge, no adverse effects were recorded in 40Fp8-vaccinated ewes, in contrast to mock-vaccinated animals that showed clear liver and reproductive organ pathology, foetal affectation and/or abortion. In some cases, Cq values above threshold were obtained by RT-qPCR in umbilical vein blood and in some foetal tissues. However, virus isolation trials in these samples was always negative. A possible explanation for the detection of viral RNA should be related with presence of residual RNA genomes derived from the challenge inoculum, since the dose administered was quite high or to the presence of viral replication intermediates since, in all cases, antibodies against the viral nucleoprotein N were detected. Failure to isolate infectious virus may also indicate that it is rapidly neutralized by vaccine-induced maternal antibodies, preventing widespread dissemination to reproductive organs or more susceptible foetal tissues.

In our experimental setting, the pregnant ewes vaccinated with 40Fp8 were sacrificed at various times post-inoculation in order to screen for potential pathological alterations in reproductive and foetal tissues. Therefore, the pregnancy was not allowed to progress to full term ending with lamb delivery. Based on the results described here it should be expected that pregnant ewes vaccinated with 40Fp8 should deliver healthy lambs after a subsequent virulent RVFV challenge. More analysis will be needed to assess the duration of protection conferred by the administration of a single dose, the effect of a booster dose or the use of immunostimulatory molecules or adjuvants to improve the protective capabilities. These experiments should be performed ideally in a natural, RVFV-endemic setting, warranting further efforts in additional 40Fp8 safety studies, such as reversion to virulence upon serial passage in animals or vector competence studies, that would impact the clinical development of 40Fp8 as a novel RVF vaccine candidate.

## Acknowledgements

We thank the animal care staff, veterinary and immunopathology services and biosafety staff at CISA for their support with the *in vivo* experiments, and Nuria de la Losa for excellent technical assistance. This work was supported by grant AGL2017-83226-R and by grant PdC2021-121717-I00 funded by MCIN/AEI /10.13039/501100011033 and by Unión Europea Next Generation EU/ PRTR.

## SUPPLEMENTARY FIGURES

**Supplemental figure 1.**
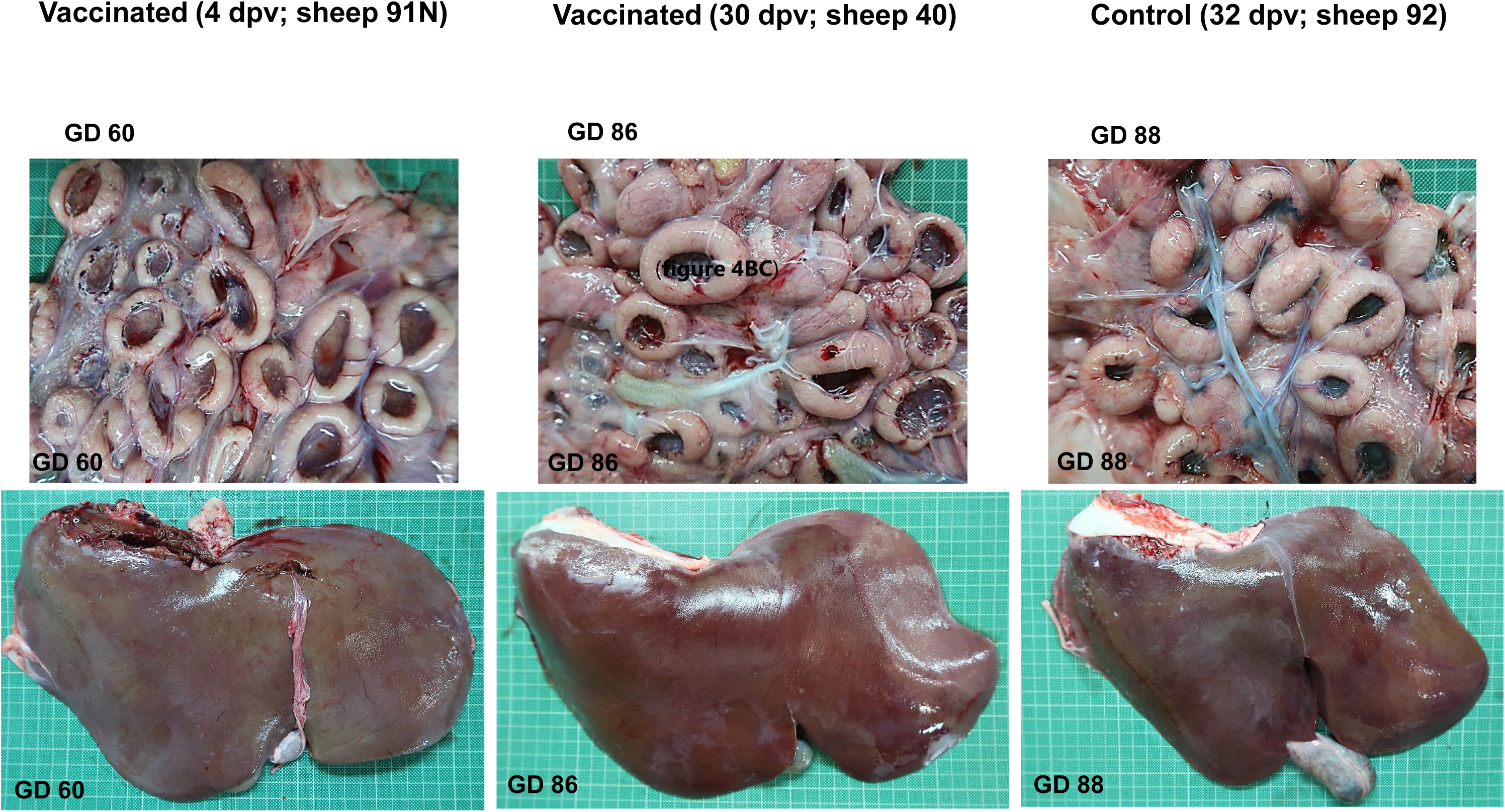
Macroscopic images of placentas and livers from ewes at different days post vaccination (dpv) with the 40Fp8 virus, corresponding to different gestational days (GD). Control: mock-inoculated ewe.

**Supplemental figure 2.**
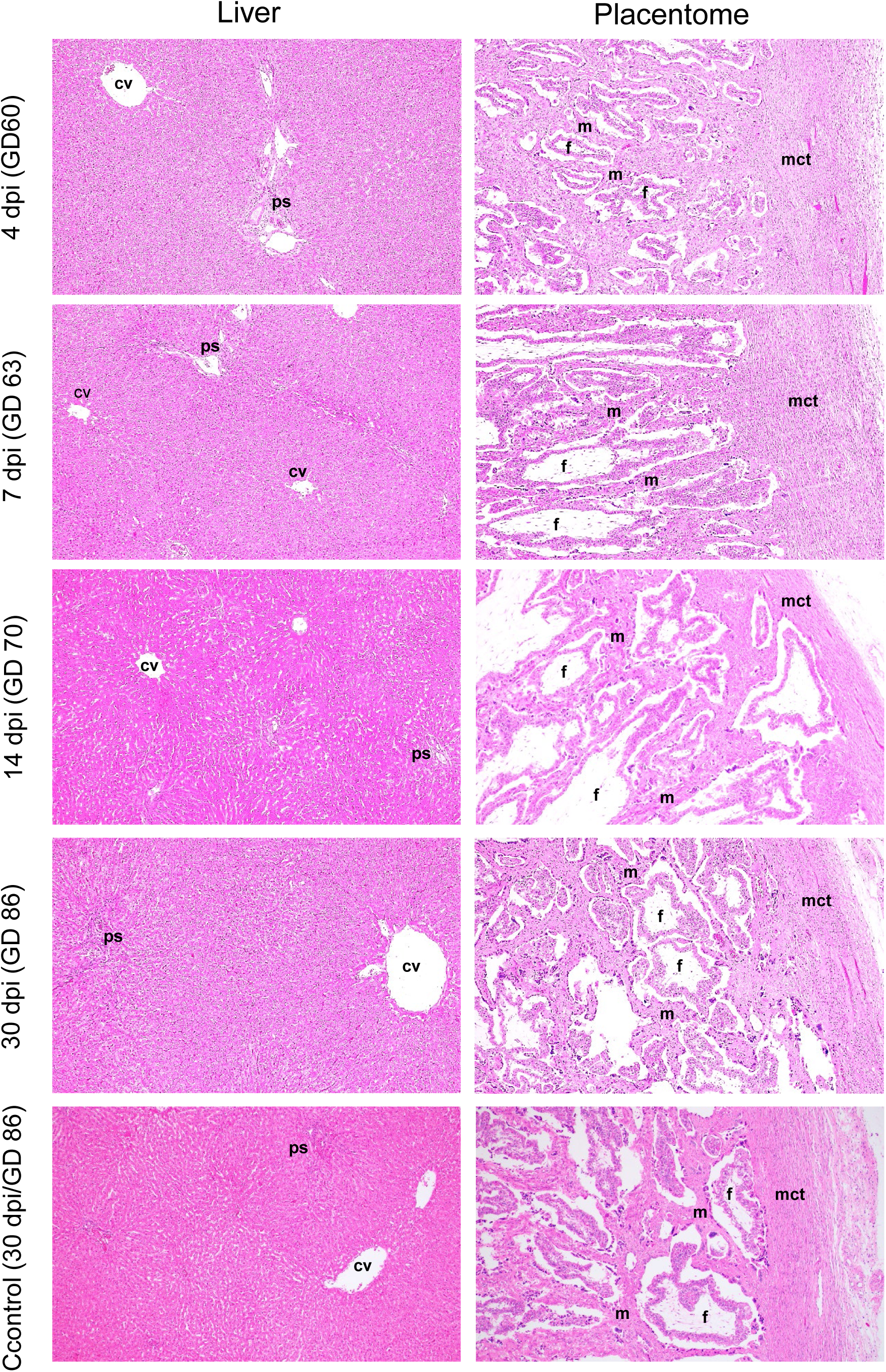
Representative histopathological images of livers and placentomes sampled from pregnant ewes subcutaneously immunized with RVFV-40Fp8 strain and euthanized at 4, 7, 14 and 30 days post-immunization (dpi). Non-immunized control pregnant ewes were euthanized at the end of the experiment (30 dpi). As observed in the non-immunized controls, neither the livers nor the placentomes of the immunized ewes showed remarkable histopathological lesions on any of the days of the experiment. Haematoxylin-eosin staining; Magnifications: Liver (4x), Placentome (10x); **(GD):** gestational day; **(cv):** centrilobular vein; **(ps):** portal space; **(f)** foetal villus; **(m)**; maternal villus; **(mct):** maternal connective tissue.

